# Primary glia cells from bank vole propagate multiple rodent-adapted scrapie prions

**DOI:** 10.1101/2021.08.06.455381

**Authors:** Karla A. Schwenke, Joo-Hee Wälzlein, Agnieszka Bauer, Achim Thomzig, Michael Beekes

## Abstract

Since the beginning prion research has been largely dependent on animal models for deciphering the disease, drug development or prion detection and quantification. Thereby, ethical as well as cost and labour-saving aspects call for alternatives *in vitro*. Cell models can replace or at least complement animal studies, but their number is still limited and the application usually restricted to certain strains and host species due to often strong transmission barriers. Bank voles promise to be an exception as they or materials prepared from them are uniquely susceptible to prions from various species *in vivo, in vitro* and in cell-free applications. Here we present a mainly astrocyte-based primary glia cell assay from bank vole, which is infectible with scrapie strains from bank vole, mouse and hamster. Stable propagation of bank vole-adapted RML, murine 22L and RML, and hamster 263K scrapie is detectable from 20 or 30 days post exposure onwards. Thereby, the infected bank vole glia cells show similar or even faster prion propagation than likewise infected glia cells of the corresponding murine or hamster hosts. We propose that our bank vole glia cell assay could be a versatile tool for studying and comparing multiple prion strains with different species backgrounds in a single cell assay.

## Introduction

Prion diseases are transmissible, fatal neurodegenerative disorders which include Creutzfeldt-Jakob disease (CJD) in humans, scrapie in sheep and goats, bovine spongiform encephalopathy (BSE) in cattle and chronic wasting disease (CWD) in cervids. They are caused by prions – proteinaceous infectious particles – which essentially consist of host-encoded, aggregated and self-propagating prion protein (PrP) [1]. The pathology is marked by abnormal PrP deposits, astrogliosis, neuronal loss and vacuolation, which lead to severe brain dysfunction and inevitably death. Despite intense research in the past decades no treatment is available so far. According to the generally accepted “protein-only” hypothesis the underlying etiological event is the conversion of the physiological, cellular prion protein PrP^C^ to its pathological isoform PrP^Sc^, which is set-off due to (i) spontaneous misfolding and aggregation, (ii) inherited genetic reasons or (iii) PrP^Sc^ acquired by infection. Once PrP^Sc^ is present, prion propagation is an autocatalytic and self-sustaining process occurring mainly in cells with high PrP^C^ expression like neurons and astrocytes [2-4].

Before gradually identifying prions and unravelling their nature, prion research was mostly dependent on animal bioassays, but even nowadays they are the gold standard for PrP^Sc^ detection and titre determination [5,6]. Refinement, reduction and replacement (“3R”) of animal studies is desirable due to ethical reasons but also towards less cost and labour-intensive methods, therefore alternatives to bioassays such as versatile cell culture models are needed. Already early in prion research a first persistently infected cell line from mouse was established [7], later further cell models were introduced (for review see [8]). One successful example of a bioassay alternative is the standard scrapie cell assay based on subcloned neuroblastoma N2a cells, which was developed for titre determination of murine scrapie (22L and RML) reaching equal sensitivity while being significantly faster and cheaper [9]. However, this assay only works for mouse-adapted scrapie strains, which is a common problem of most cell culture models: being limited to certain host-adapted prions only [10,8].

While neurons are extensively studied in prion disease, the role of other cell types remain to be further elucidated. Although prion accumulation leads to neuronal degeneration or death, also non-neuronal cells such as astrocytes are infected and potent propagators of prions *in vivo* but also *in vitro* [11]. Nonetheless, cell models mainly exist for neurons with few exceptions for primary hamster [12] or mouse [13,11] glia cells, human stem-cell derived astrocytes [14] or recently the first infectible astrocyte cell line [15]. Due to transmission barriers hamster or murine cells are generally restricted to infection with species-adapted prion strains. Bank voles, however, were found to be universal acceptors for prions [16] making them fast *in vivo* models for CJD [17] as well as CWD [18], but also broaden the application area of *in vitro* assays like real-time quaking induced conversion (RT-QuIC) [19,20] and protein misfolding cyclic amplification (PMCA) [21]. So far only two cell lines expressing bank vole PrP were engineered being susceptible to murine scrapie (22L) and native CWD [22], but no primary bank vole cell culture infectible with prions exists to the best of our knowledge. Cell lines have the advantage of providing fast and cheap cell assays, which are easy to maintain and have the potential for gene-editing. Benefits of primary compared to immortalised cells include a more natural simulation of an infection; for instance persistently prion-infected cell lines do not show cytotoxicity in contrast to prion-infected cells *in vivo* [13,23]. Also, in case of primary glia cells the post-mitotic state allows long-term cultivation without the need for passaging, which mimics the long incubation periods of prions and slow PrP^Sc^ accumulation *in vivo* and further eliminates the risk of losing infectious titres due to disproportionally faster cell proliferation than prion propagation [24]. A primary bank vole cell assay infectible with prions could provide a tool for investigating a variety of prion strains from different host species under more natural conditions in a single cell model. In analogy to our already existing, robust hamster glia cell assay [12,25], we aimed to establish a primary glia cell culture from bank vole, which would be susceptible to multiple prion strains from different species and exhibit stable prion propagation. Initially, a proof-of-concept with multiple rodent-adapted scrapie strains was to be accomplished, in order to provide a platform for the testing of more strains with different species background and native isolates such as CWD or CJD in the future. Here we present a primary mainly astrocyte-based cell assay from bank vole, which shows stable propagation of four scrapie strains from bank vole, mouse and hamster.

## Materials and Methods

### Animals and tissues

Neonatal C57BL/6 mice and bank voles (*Myodes glareolus*, PrP genotype Bv109M [18]) were obtained from colonies in the animal facility of Robert Koch Institute (RKI). Initial breeding pairs of bank voles were received from Istituto Superiore di Sanità, Rome, Italy.

Brain tissue of terminally ill hamsters with 263K were taken from stock at RKI. Murine 22L, RML and ME7 as well as bank-vole RML and ME7 brain tissue samples were kindly provided by the Gilch and Schätzl Lab, University of Calgary, Canada.

### Glia cell isolation and cultivation

Primary glia cells were isolated from brains of neonatal bank voles (0-3 days post natum) based on Buonfiglioli *et al*. [26] with modifications. Briefly, the cerebellum, olfactory bulb and meninges were removed and the brains washed three times with ice-cold Hank’s Balanced Salt Solution (HBSS). Enzymatic dissociation was carried out by incubating brain tissue with 10 mg/ml Trypsin (Biochrom, Germany) and 0.5 mg/ml DNase I (Roche, Switzerland) in HBSS for two minutes at first and subsequently with 5 mg/ml DNase I in HBSS while gently triturating by pipetting. The tissue suspension was centrifuged at 120xg for 10 min, cells resuspended in neurobasal medium containing 1% G5 supplement, 1% GlutaMAX and 100 U/ml Penicillin/Streptomycin (all Gibco™ Thermo Fisher, USA; referred to as NBM complete) and quickly plated on poly-L-lysine (PLL; 0.01% (w/v); Sigma-Aldrich, USA)-coated T75 flasks, so that 2.5 brains were allocated per flask. Cells were maintained under humid conditions with 5% CO_2_ at 37 °C. After two days some adherent cells but mainly semi-adherent 3D cell aggregates could be observed. In order to reduce cytotoxic cell debris and tissue chunks, cells were washed once very gently with phosphate-buffered saline (PBS) and incubated for further two days. During this time most cells became adherent and could be washed thoroughly for removal of remaining cell debris. After another two to three days cells have grown confluent and were split in a ratio of 1:3. After further seven days (14 days post isolation) cells were ready to be harvested and either directly seeded for experiments or cryo-preserved for later use.

Glia cells from neonatal C57BL/6 mice were isolated likewise, except that Dulbecco’s Modified Eagle’s Medium (DMEM; Gibco™ Thermo Fisher, USA) containing 10% foetal bovine serum (FBS; Pan Biotech, Germany, P30-3031), 1% GlutaMAX and 100 U/ml Penicillin/Streptomycin was used instead of NBM complete. Due to the presence of serum cells were adherent after two days post isolation already, could be washed thoroughly for four times and were cultivated for another five days until cells have grown confluent. Seven days post isolation mouse glia cells were harvested and used for experiments without further passaging.

### Flow cytometry

For determining the cellular composition of the bank vole glia culture at the time of infection, expression of the astrocyte-specific markers glial fibrillary acidic protein (GFAP) and glutamate transporter 1 (GLT-1) was investigated as well as PrP expression. Samples were analysed in triplicates and the corresponding Fluorescence Minus One (FMO) controls in duplicates. Furthermore, unstained and compensation controls were performed for necessary adjustments of the laser channels and compensation of fluorophores. Briefly, glia cells of a confluent T75 flask were detached with Accutase (Capricorn Scientific, Germany), centrifuged, resuspended in FACS buffer (PBS containing 2% (v/v) FBS and 0.1% (w/v) sodium azide) and equally distributed on a round-bottom 96-well plate. Prior to intracellular staining cells were fixed with 2% (w/v) paraformaldehyde, cell membranes permeabilised with 0.5% (w/v) saponin and unspecific binding sites blocked with 1% (w/v) bovine serum albumin. Cells were then incubated for 30 min on ice with previously titrated dilutions of primary antibodies [recombinant anti-GFAP PE-conjugated (REA335, Miltenyi Biotec, Germany): 1:100; polyclonal rabbit anti-GLT-1 (ab41621, Abcam, UK): 1:200; monoclonal mouse anti-PrP (SAF84, Bertin, France): 1:400] and subsequently, after washing once with FACS buffer, for 15 min at room temperature with the secondary antibodies [Alexa488-conjugated anti-rabbit IgG and Alexa647-conjugated anti-mouse IgG: 1:500 (Thermo Fisher, USA)]. Afterwards cells were washed twice and finally resuspended in 200 µl FACS buffer. 150 µl of each sample was analysed using the MACSQuant Analyzer 10 (Miltenyi Biotec, Germany) and the data evaluated using FlowJo 10 software (BD Life Science, USA). Cell debris and doublet cells were excluded based on scatter signals.

### Protein misfolding cyclic amplification

In order to test *in vitro* engineered PrP^Sc^ seeds for infectivity protein misfolding cyclic amplification (PMCA) was performed of the strains BV-RML or hamster 263K in a bank vole brain substrate as previously described [27] with following parameters. 90 µl of PMCA substrate [10% (w/v) bank vole brain homogenate in conversion buffer: PBS, 1% (v/v) Triton X-100, 6 mM EDTA, 100 mM NaCl, protease-inhibitor cocktail cOmplete (Roche, Switzerland), pH 7.4] and two Teflon beads (1/16”; McMaster-Carr, USA) were added to each micro reaction tube prior to adding 10 µl of the respective seed (10^−2^ diluted 10% (w/v) brain homogenate of BV-RML or 263K or normal bank vole as negative control). Four rounds consisting of 48 cycles each were conducted with sonication of 170 W power for 30 s every 29.5 min at 37 °C in a Q700 microplate horn sonicator (QSonica, USA). Samples were passaged 1:5 to the next round and mixed with 80 µl fresh substrate.

Prior to using the PMCA products of the fourth round as infectious seeds in cell culture, the conversion buffer with its cytotoxic components was substituted with PBS. For this purpose, PMCA products were centrifuged at 45,000 rpm for 2.5 h at 4 °C (Optima™ Max Ultracentrifuge, Beckman Coulter, USA), the supernatants discarded and the pellets resuspended in an equal volume of PBS.

### Cell-based infection assay

For the infection assay medium for mouse glia cells was changed from DMEM to DMEM/F12 (Gibco^™^ Thermo Fisher, USA) with the same supplements for better cell viability during long-term cultivation, and for bank vole cells NBM complete was switched to DMEM complete supplemented with 1% N-2 supplement (Gibco^™^ Thermo Fisher, USA). Mouse or bank vole glia cells were plated at 1×10^5^ or 5×10^4^ cells/well, respectively, in PLL-coated 6-well plates and grown semi-confluent for five days. Seeds as either 10% brain homogenates diluted 1:10 in the respective cell culture medium or purified PMCA products were always sonicated for 30 s at 300 W in a microplate horn sonicator (Misonix 3000, USA) prior inoculation. Cells were exposed to 10 µl of the respective seeds in 2 ml medium/well for three days, subsequently washed once with PBS for removal of non-incorporated seeds and supplemented with 3 ml fresh medium. Cells were cultivated until the indicated time points with a weekly medium change. For all experiments one sample was taken at 3 dpe as a reference for initial seeds, further cells were harvested at 20, 30, 40, 45 or 60 dpe as stated by lysing cells with 100 µl/well of 1% sarcosyl.

### SDS-PAGE and Western blot

Cell lysates in 1% sarcosyl were digested with 75 µg/ml proteinase K (PK; Roche, Switzerland) for 1 h at 37 °C and subsequently denatured in an equal volume of 2x sample loading buffer at 110 °C. Deglycosylation was performed with peptide-*N*-glycosidase F (PNGase F; New England Biolabs, USA) following the manufacturer’s instructions [28]. For this 15 µl PK-digested and denatured samples were mixed with 2 µl each of 10x GlycoBuffer2, NP-40 and PNGase F and incubated for 2 h at 37 °C. 20 µl of either type of samples were run on BOLT 4-12% Bis-Tris mini protein gels for SDS-PAGE and electroblotted using the *iBlot 2* dry blotting system (Thermo Fisher, USA). Polyvinylidene fluoride membranes were probed with anti-PrP monoclonal antibody ICSM-18 (1:4,000; D-Gen, UK) overnight and anti-mouse IgG alkaline phosphatase-linked secondary antibody (1:5,000; Dako, USA). Membranes were incubated with CDP-Star chemiluminescent substrate (Thermo Fisher, USA) for alkaline phosphatase chemiluminescence reaction and detected on *Amersham Hyperfilm*^*TM*^ ECL films (GE Healthcare, USA). Images were created in Illustrator 2021 (Adobe, USA).

## Results

### Establishment of primary bank vole glia cell culture

The primary bank vole glia cell culture (BV-Glia) was established based on our protocol for Syrian hamster glia cells [12,25] and the protocol for murine C57BL/6 glia cells (BL6-Glia) from the Kettenmann Lab (Max Delbrück Center, Berlin, Germany) with modifications specific for bank vole glia. The conventional protocol of isolating cells directly into serum-containing DMEM resulted in a very strong proliferation of bank vole cells, making them inapplicable for infection assays with required long-term cultivation without passaging. Changing the isolation protocol to a serum-free neurobasal medium supplemented with astrocytic-growth-promoting Bottenstein’s G5 supplement first induced an intermediate state of 3D cell aggregate formation during the first days until cells differentiated and became adherent with a distinct fibrous morphology, which is similarly described in the literature [29,30]. Only after obtaining glia cells in a more mature state towards glial differentiation in the second passage, cells were used for infection assays and culture medium was switched to serum-containing DMEM to stabilise cells.

### Characterisation of the glia cell culture

Since no specific astrocyte selection was performed, composition of the mixed glia cell culture was investigated by flow cytometry at the same passage and time point after plating cells analogue to prion inoculation. For characterisation the common astrocyte-specific cell markers glial fibrillary acidic protein (GFAP) and glutamate transporter 1 (GLT-1) were used. Additionally, expression of PrP^C^ was investigated since this is known to be essential for sufficient prion propagation [31,32]. Flow cytometry analysis (figure 1) revealed a high content of astrocytes in the mixed culture with 94% in the double-positive GFAP+ GLT-1+ cell population after gating for intact and single cells, of which more than 92% were GFAP+ and almost 97% GLT-1+. These cells showed an overall high PrP expression on >99% of the cells. Further consistence of the remaining negative cell population was not determined.

**Figure 1:**
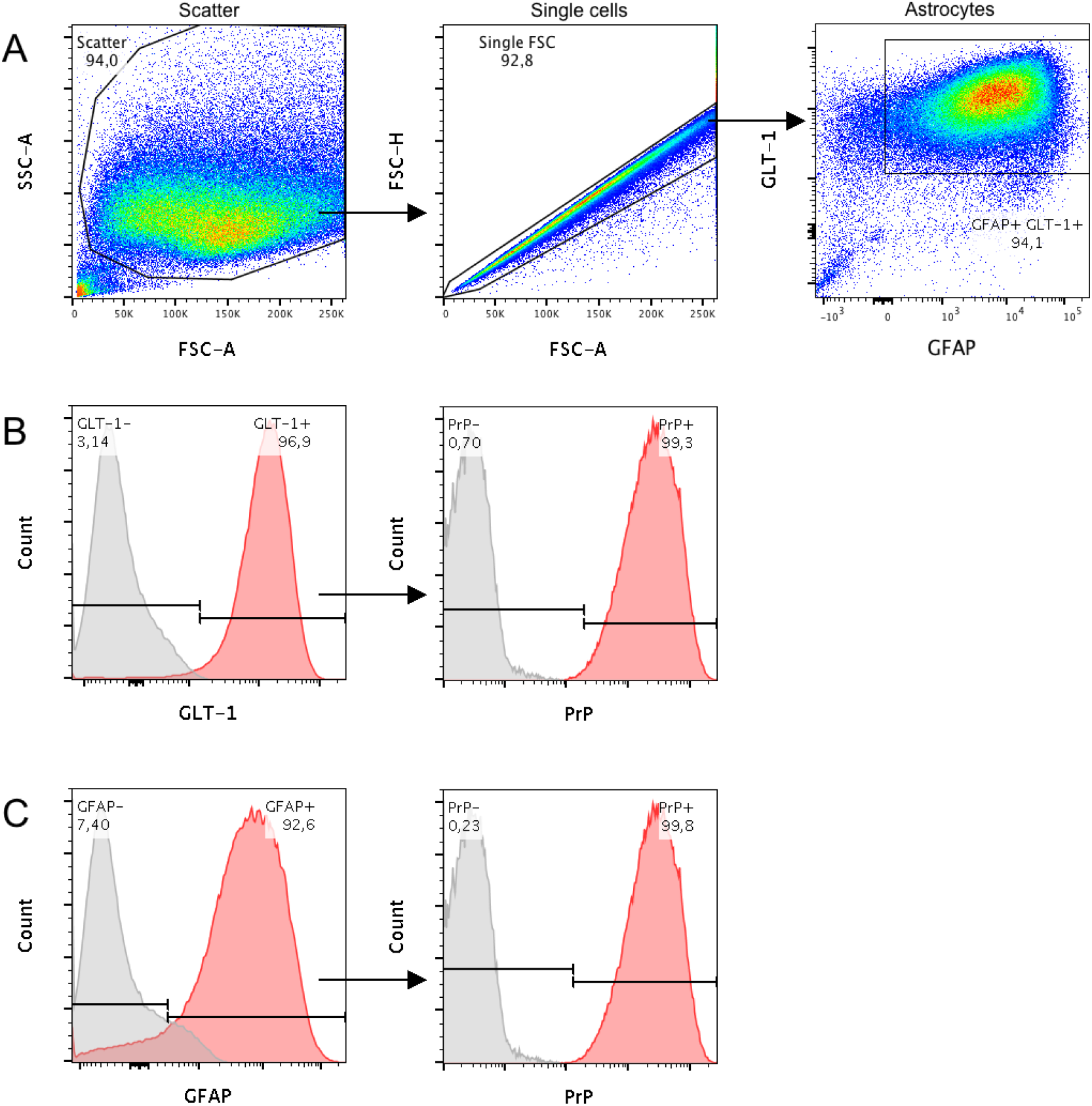
Composition of primary bank vole glia cells at the time of infection. (20 days post isolation, passage 2). Characterisation was performed by flow cytometry analysis based on expression of GFAP and GLT-1, two specific markers for cells of astrocytic phenotype. Additionally, levels of PrP^C^ expression were determined as crucial factor for prion propagation. (A) Proportion of the analysed cells, which were found to co-express GFAP and GLT-1. Cell debris and doublets were excluded from the analysis by gating as illustrated. (B) Histogram of those cells, which were stained for GLT-1 and (C) for GFAP. PrP^C^ expression of each subpopulation is shown in the respective second panel. Red areas of the histograms refer to stained samples and grey areas to the corresponding control incubated without the target antibody. The experiment was carried out in triplicate, of which one representative is shown. According to the analysis the glia cell culture mainly consisted of astrocytes with high PrP^C^ expression, which provided the basis for an infectible cell assay. FSC: forward scatter, SSC: side scatter

### PrP^Sc^ propagation in the glia infection assay

BV-Glia were inoculated with rodent-adapted scrapie prions 22L and RML from C57BL/6 mice (BL6-22L, BL6-RML; n=3), bank vole-adapted RML (BV-RML; n=3) and 263K from hamster (n=2). For each inoculum the corresponding, non-infectious brain homogenate (NBH) was used as mock control. Cells were exposed to each inoculum (10 µl of 1% brain homogenate) for three days, subsequently unattached material was removed and first samples were harvested as reference for residual inoculum (referred to as 3 dpe). All four scrapie strains were efficiently propagated in BV-Glia, which was detected via western blot (figure 2). Thereby, clearly increasing amounts of protease-resistant pathological PrP (PrP^res^) could be observed starting with first faint signals at 20 dpe and manifesting at 30 dpe onwards (BL6-22L, BL6-RML, BV-RML) or distinct signals already at the earliest time point at 20 dpe, respectively (263K). The increase of PrP^res^ from the reference time point at 3 dpe to the following time points at 20 to 60 dpe indicates accumulating PrP^Sc^ formation. Mock-infected cells were always found to be negative for PrP^res^ in Western blot (not shown).

**Figure 2:**
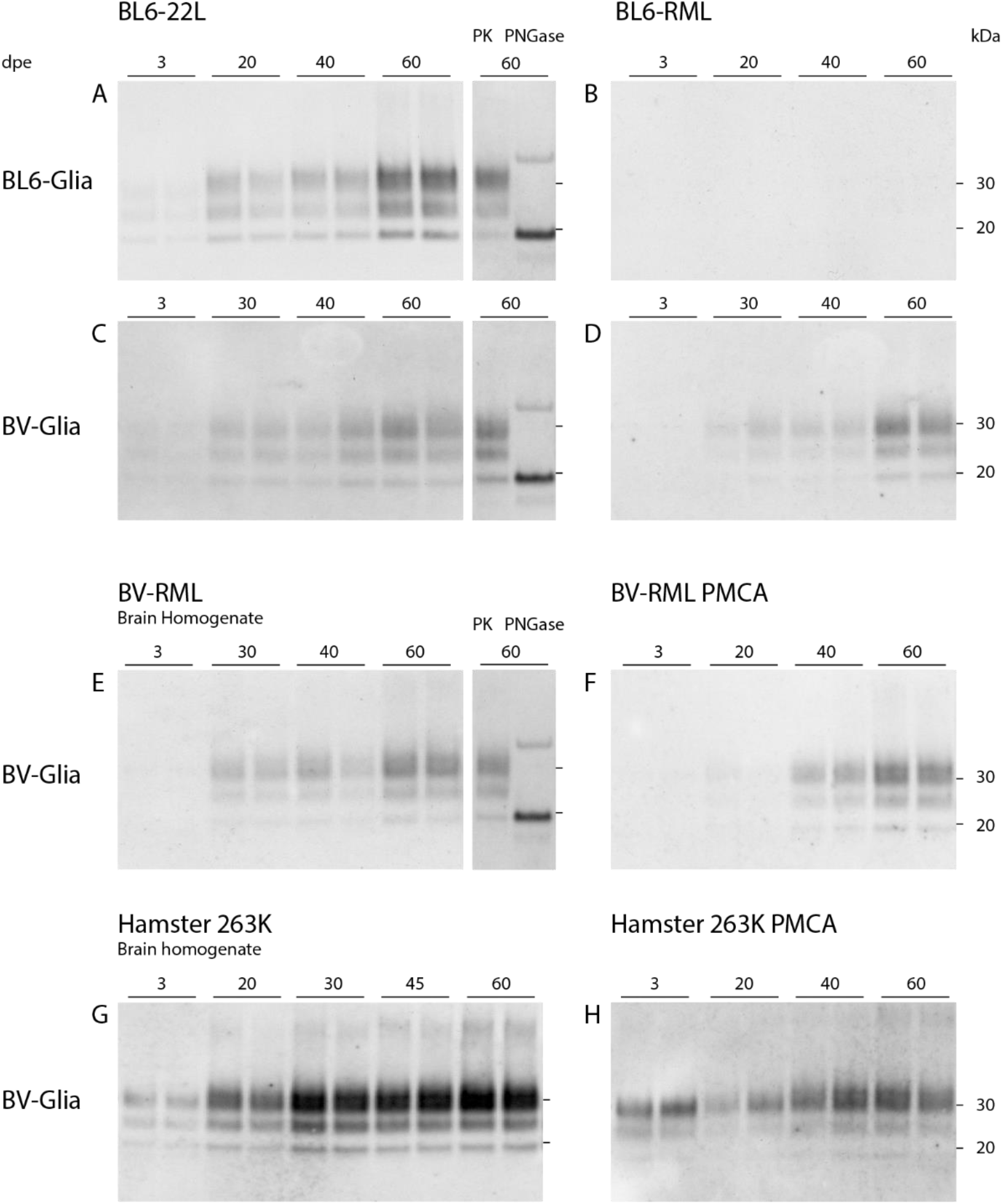
Western blot analyses of PK-digested cell lysates of prion-infected glia cells from C57BL/6 mice and bank vole. 1×10^5^ cells/well from mouse (A, B) or 5×10^4^ cells/well from bank vole (C-H) were exposed to scrapie prion strains: mouse-adapted 22L (A, C), mouse-adapted RML (B, D), bank vole-adapted RML (E) and hamster-adapted 263K (G) in a concentration of 10 µl of 1% brain homogenate or 10 µl of bank vole PMCA product of BV-RML (F) or 263K (H) for three days and harvested at the indicated time points. The immunoblots illustrate the propagation of pathological prion protein in cells over the incubation time dependent on the seeding material. All four strains and both PMCA products show an efficient propagation in bank vole glia cells, even for BL6-RML which shows no sign of infection in mouse cells up to 60 dpe. In A, C and E 60 dpe-samples deglycosylated with peptide-N-glycosidase F (PNGase F) are shown with a clear shift of the di-and monoglycosylated band to the unglycosylated band at 19 kDa. The band at ∼35 kDa is considered an artefact of the deglycosylation process itself [28]. Representative Western blots of infection assays are shown, which were stained with anti-PrP antibody ICSM-18 (1:4,000) (n=3, except: 263K: n=2, BV-RML-PMCA: n=2 and 263K-PMCA: n=1; all in duplicates).

BL6-22L-infected BL6-Glia (n=3) showed a slightly faster and stronger prion propagation compared to the bank vole cells (distinct Western blot signals already at 20 dpe compared to 30 dpe, figure 2 A). In contrast, inoculation with BL6-RML (n=3) revealed a less efficient prion propagation in BL6-Glia, for which in only one of three experiments a very faint signal could be seen at 60 dpe (figure 2 B). This indicates infectibility in principle but need for a prolonged incubation time. Surprisingly, other than the murine cells PrP^res^ signals in BL6-RML-inoculated BV-Glia were detectable already from 30 dpe onwards (figure 2 D), comparable to BL6-22L or BV-RML prions in bank vole cells.

Also, inoculation of BV-Glia with hamster 263K revealed an efficient prion propagation with strong signals as early as 20 dpe already reaching maximal propagation only 10 days later, which is on the same level as known from the currently established version of our hamster glia cell assay (unpublished data).

In contrast, mouse and bank vole-adapted ME7 scrapie brain homogenates were tested twice under the same conditions but did not show any sign of propagation up to 60 dpe in neither BL6 nor BV-Glia (not shown).

In order to verify the Western blot signals as PrP^res^, deglycosylation was carried out with the N-glycosidase PNGase F. For the representatively tested cell lysates (cells infected with BL6-22L and BV-RML at 60 dpe) a uniform shift of the di-and monoglycosylated bands to the unglycosylated band at approximately 19 kDa could be observed as expected (Figure 2 A, C, E), which confirms the identity of the immunolabelled prion protein.

Additionally, infectibility of BV-Glia with PMCA products of BV-RML and 263K was investigated in a pilot test. BV-RML and 263K brain homogenates were subjected to PMCA for four rounds each in a bank vole brain substrate (not shown). Subsequently, cells were inoculated the same way as previously described with 10 µl of undiluted purified PMCA product or the respective PMCA negative control without seeds as mock control. Pilot results show replication for both PMCA seeds requiring only a few more days for reaching the same levels of propagated PrP^res^ as their original brain homogenates (figure 2 F+H as compared to figure 2 E+G, respectively), while mock-infected cells did not show any PrP^res^ (not shown). These results need to be reproduced for confirmation of their robustness and a more detailed comparison between PMCA-adapted and original seeds.

## Discussion

### A mainly astrocytic primary bank vole glia assay suitable for long-term cultivation

To the best of our knowledge this is the first report of a primary glia cell culture from bank vole, which shows a stable propagation of rodent-adapted scrapie prions of murine, hamster and bank vole origin. The primary mixed glia cell culture revealed a high content of astrocytic cells in flow cytometry, of which 94% belonged to the double-positive cell population for the specific astrocyte markers GFAP and GLT-1. While GFAP has been seen as a definite single astrocyte marker for a long time, nowadays this assumption is revisited and the use of at least two markers recommended [33]. We included GLT-1 in our analysis, which is a potent marker for cells with astrocytic phenotype [34] and the predominant glutamate transporter in the developed brain [35,36]. Interestingly, in a first flow cytometry analysis for test purposes a tremendous increase of GLT-1+ cells was observed between the initially applied protocol with serum-containing medium and the subsequently adapted bank vole protocol with serum-free cultivation for cell isolation (not shown), which points to more mature astrocytes in the cell mix of the latter. Previous studies [37,38] have shown that exposing freshly isolated glia cells to serum changes their expression profile and apparently keeps them in an immature state as opposed to serum-free cultivation. In contrast, a later cultivation in serum does not seem to alter cell composition anymore. This would explain the impeded proliferation of the bank vole glia cells and the quicker shift to a post-mitotic state when the isolation protocol was changed. It remains unclear why absence of serum does not appear to be crucial for mouse glia cells, but possibly efficiency of the BL6-Glia infection assay could be further enhanced by switching to serum-free cell isolation. The combination of this with the astrocytic-growth-promoting Bottenstein’s G5 supplement likely contributes to the relatively pure astrocyte culture with little contamination of other cells like microglia or oligodendrocytes, which are not considered a site of pronounced prion propagation [39-41]. Hence, this isolation protocol seems to promote the differentiation of the cells to a more mature astrocyte cell type, which seems to favour prion propagation.

### Equal or faster propagation rate of scrapie prions in bank vole glia compared to murine cells

In this bank vole glia cell assay all four scrapie strains led to a productive infection and PrP^Sc^ accumulation within relatively short incubation periods with immunoblot signals being first visible at 20 or latest 30 dpe. Thereby, the murine strains showed an almost similar fast propagation (BL6-22L) as or even an accelerated one (BL6-RML) within half of the incubation time than in likewise infected BL6-Glia. Even for the non-murine 263K hamster scrapie strain the propagation rate was as fast as in hamster glia cell assays carried out in our laboratory. This points to a broader and increased susceptibility of prion strains in bank vole cells as already reported *in vivo* and *in vitro* and suggests a highly versatile usability of this glia-cell-based assay.

The ME7 scrapie strain is known to be slower in propagation *in vivo* and in cell culture, and to show a different tropism than e.g. 22L or RML [42-44,23]. Since ME7 was found to propagate in neurons, it seems possible that this mainly astrocyte-based assay is refractory to ME7 infection in general or at least needs substantially prolonged incubation periods. A complementary neuronal cell assay from bank vole could possibly provide more insights into the tropism of ME7.

Our pilot experiments with PMCA products as seeding-active materials demonstrate an infectibility of the BV-Glia with such *in vitro* generated seeds as well. The presumably less productive infection with PMCA products compared with the original brain-derived seeds has already been shown in *in vivo* and *in vitro* infection models [45,46], but more experiments would be necessary to confirm our results and compare both types of seeds in more detail. The amount of seeds in the 263K PMCA product appeared relatively high according to the strong signal at 3 dpe, but the declining signal at 20 dpe at first and later clearly increasing signal from 40 dpe on and including a slight shift in the diglycosylated band points to non-residual PrP^Sc^ accumulated by glia cells. For prion-infected cell lines analyses of PrP^Sc^ formation is generally carried out after (serial) passaging of cells to exclude residual inoculum. Since no cell passaging was performed after infecting our primary cells, we consider the clear increase of PrP^res^ over time in Western blot as evidence for biological PrP^Sc^ amplification in our bank vole glia cells.

### Bank vole as universal prion acceptor in a cell model

Our results further confirm the assumption that bank vole is a universal acceptor for prions in cell culture as well. Remarkably, not only the transmission barrier is lower, but partly also the incubation time even shorter than for murine prions in mouse cells. This assay therefore promises to provide a versatile tool covering a range of prion strains from different host species, which could be used for studying prion infections as well as performing drug development or decontamination studies. Thereby, this cell assay has the potential to provide a single, universal platform, which could combine studies with several prion strains in one assay and improve their comparability in-between. With the exception of stem cell-derived astrocyte or cerebral organoid cultures [14,47], no further cell model propagating human CJD prions exists to date. Bank voles with their exceptional susceptibility for various prion strains raise expectations for providing broad cell models potentially infectible with CJD. Furthermore, our cell assay is cheaper, easier and less labour-intensive than stem-cell-derived astrocyte or organoid cultures, and notably bioassays. Beyond the aim of implementing the 3R principles for the sake of animal protection in general, bank voles demand particular attention in this direction since they are unusually prone to stress and require extremely careful handling [27]. Replacing animal experiments with the use of bank vole-tissue for *in vitro* studies would therefore represent a highly desirable step forward in terms of animal welfare.

## Declarations

### Ethical statement

According to German regulations euthanasia of normal wild-type animals for experimental purposes does not require approval by ethics committees. However, sacrificed animals were voluntarily reported to the Authority for Animal Protection in Berlin (“Landesamt für Gesundheit und Soziales Berlin”, Berlin, Germany; T 0300/15, TN 0001/21 and T 0256/15). All animals used in this study were handled under the European directive regarding the protection of animals used for scientific purposes in strict accordance with the German Animal Welfare Act (Tierschutzgesetz).

### Competing interests

The authors declare that they have no competing interests.

### Funding

KAS was financially supported by Alberta Prion Research Institute (Alberta, Canada, project number PEX12015) and RKI. JHW was financially supported by German Federal Ministry of Education and Research (funding initiative „Alternative methods to animal experiments”, project number 031L0065).

### Authors’ contributions

KAS and JHW conceived and performed the experiments and analysed the results. AB guided the flow cytometry experiment and analysed the data. KAS, JHW, AT and MB contributed to study conceptualisation and data interpretation. MB raised the funding. KAS drafted the manuscript. All authors revised and approved the final manuscript.

## Acknowledgments

We are grateful to Helmut Kettenmann (Max Delbrück Center, Berlin, Germany) for sharing his protocol for murine primary glia cell isolation and to Maren Wendt for the practical training; to Julia Madela-Mönchinger (RKI) for her flow cytometry guidance; to Umberto Agrimi and Michele Di Bari (Istituto Superiore di Sanità, Rome, Italy) for kindly providing breeding pairs of bank voles; to Sabine Gilch and Hermann Schätzl (University of Calgary, Canada) for kindly providing the murine and bank vole-adapted scrapie strains and to the animal facility of RKI for their support.

## Notes

### Competing Interest Statement

The authors have declared no competing interest.

